# Target-driven design of a coumarinyl chalcone scaffold based novel EF2 Kinase inhibitor suppresses breast cancer growth *in vivo*

**DOI:** 10.1101/2020.11.06.371062

**Authors:** Ferah Comert Onder, Nermin Kahraman, Esen Bellur Atici, Ali Cagir, Hakan Kandemir, Gizem Tatar, Tugba Taskin Tok, Bekir Karliga, Serdar Durdagi, Mehmet Ay, Bulent Ozpolat

## Abstract

Eukaryotic elongation factor 2 kinase (eEF-2K), an unusual alpha kinase, is involved in protein synthesis through phosphorylation of elongation factor 2 (EF2). eEF-2K is indicated as one of the critical drivers of breast cancer and associated with poor clinical prognosis, representing a potential molecular target. The crystal structure of eEF-2K is unknown and there is no potent and effective eEF-2K inhibitor reported for clinical applications. To meet this challenge, we designed and synthesized several generations of potential inhibitor compounds and performed *in silico* studies. The effect of the inhibitors at the binding pocket of eEF-2K is analyzed after developing a 3D target model by homology modeling approaches using a domain of another α-kinase called myosin heavy-chain kinase A (MHCKA) that is closely resembling eEF-2K. Our results showed that compounds with coumarin-chalcone cores have a high predicted binding affinity for binding to eEF-2K. Following *in vitro* studies, we identified a compound that was highly effective in inhibiting eEF-2K activity at submicromolar concentrations and inhibited proliferation of various breast cancer cells with different features (BT20, MDA-MB-231, MDA-MB-436 and MCF-7) by induction of apoptosis while sparing normal cells. *In vivo* systemic administration of the the lead inhibitor encapsulated in single lipid-based nanoparticles twice a week significantly supressed growth of MDA-MB-231 tumors in orthotopic breast cancer models in nude mice. In conclusion, our study provides the first *in vivo* effective small molecule eEF-2K inhibitor that may be used for molecularly targeted precison medicine strategies in breast cancer or other eEF-2K-dependent tumors.

## Introduction

Eukaryotic elongation factor 2 kinase (eEF-2K) belongs to alpha kinase family and is involved in regulation of protein synthesis by phosphorylating elongation factor 2 (EF2) as an unusual kinase^1-3^. eEF-2K has a limited sequence identity to more than 500 conventional protein kinases and cannot be inhibited by pankinase inhibitors such as staurosporin. eEF-2K is activated by calcium-calmodulin system and various oncogenic signaling, mitogens and growth factors (i.e., EGFR). Cellular and metabolic stress such as hypoxia, nutrient-deprivation and autophagy induce eEF-2K activity, suggesting that this kinase acts as a survival pathway^4-10^. eEF-2K plays a role in regulating the Warburg effect in tumor cells by modulating the synthesis of protein phosphatase 2A (PP2A) and promotes expression of c-Myc pathway pyruvate kinase (PK) M2 isoform, the key glycolytic enzyme transcriptionally activated by c-Myc^11^. Recently, adenosine monophosphate-activated kinase (AMPK), one of the key regulators of energy homeostasis, was identified as another substrate of eEF-2K^12^, revealing that eEF-2K signaling is involved in regulation of the metabolism and cellular energy.

We previously demonstrated that eEF-2K is highly upregulated in triple negative breast cancer (TNBC), BRCA1 mutated and ER+ breast cancer, and its expression is associated with poor clinical outcome, metastatic disease and shorter patient survival in TNBC^13-15^. Inhibition of eEF-2K by genetic methods blocked cancer cell proliferation, migration, invasion, and tumorigenesis in TNBC, pancreatic and lung cancer models^13-19^. More importantly, *in vivo* therapeutic inhibition of EF2K by gene-targeted therapies using RNAi suppressed growth of human cancer xenografts in various tumor models in mice, including TNBC, BRCA1-mutated breast cancer, and lung cancer^13,14,17,19^. Furthermore, it was shown that *in vivo* targeting of eEF-2K enhances the efficacy of chemotherapy in tumor models^13^. Overeall, the studies indicate that eEF-2K is one of the critical drivers of TNBC, pancreatic and lung cancers and serve as a potential molecular target. Therefore, recently, interest in developing effective inhibitors targeting eEF-2K has increased significantly^20-28^.

Several inhibitors of eEF-2K have been described in the literature^20-28^. However, these inhibitors have been neither potent nor specific for clinical appplications. For instance, 1-benzyl-3-cetyl-2-methylimidazolium iodide (NH125), an imidazolium derivative, was the first inhibitor published. Studies showed that it does not inhibit eEF2 phosphorylation when exposed to cells^29,30^. Later, A-484954 (7-amino-1-cyclopropyl-3-ethyl-2,4-dioxo-1,2,3,4-tetrahydro pyrido[2,3-d]pyrimidine-6-carboxamide), a pyrido-pyrimidinedione, was identified by Abbott laboratories as a specific eEF-2K inhibitor, but it is a weak inhibitor and inhibits cell proliferation at very high doses (IC_50_ ∼ 50-75 µM)^31,32^. Another well known and commercial available inhibitor, TX-1918, (2-((3,5-dimethyl-4-hydroxyphenyl)-methylene)-4-cyclopentene-1,3-dione) and DFTD (2,6-Diamino-4-(2-fluorophenyl)-4H-triopyran-3,5-dicarbonitrile) have been reported as eEF-2K inhibitor with IC_50_ of 0.44-1 μM and 60 μM, respectively. However, TX-1918 has been shown to inhibit also many kinases including Src, PKA, PKC and EGFR^33,34^. Thus, development of highly effective and selective inhibitors targeting eEF-2K is urgently needed for clinical translation.

The major obstacle for rationale drug design and development of effective eEF-2K inhibitors is lack of information regarding 3D structure of the catalytic domain of this kinase. To overcome this challange, we designed and screened different scaffolds of compounds and evaluated their activity by *in silico* using homology modeled target structure of eEF-2K and *in vitro* assays. Our results showed that compounds including coumarin-chalcone cores have high predicted binding affinity for bining to eEF-2K. Thus, a series of compounds including coumarin-chalcone compounds were synthesized, and using the homology modeling, compound **2C** was identified as the lead small molecule inhibitor. Further *in silico* studies (induced fit docking -IFD, quantum polarized ligand docking, molecular dynamics (MD) simulations and post-processing analysis of MD trajectories) were performed and the effect of **2C** at the binding pocket of eEF-2K was investigated in atomic details. Our *in vitro* results indicated that the lead inhibitor compound **2C** was highly effective in inhibiting eEF-2K in breast cancer cells at submicromolar concentrations (<1 µM) and showed significant antiproliferative effects by inducing apoptosis. More importantly, compound **2C** was also highly effective in inhibiting tumor growth of highly aggressive TNBC tumors *in vivo* in mice with no observed toxicity, suggesting that the **2C** may be considered for preclinical development for clinical translation to phase I clinical trials.

## RESULTS

### Homology modeling, Molecular Docking and Molecular Dynamics (MD) Simulations

Currently the crystal and 3D structure of eEF-2K is unknown. However, the three-dimensional structures of the catalytic domain of another α-kinase called myosin heavy-chain kinase A (MHCKA) is available and has been studied in considerable detail^35^. Therefore, we developed a predicted model of human eEF-2K used as a myosin heavy-chain kinase A (MHCK-A) template with homology modeling studies^36^. Next, we designed and screened different scaffolds of compounds including coumarin-chalcone scaffold and evaluated their activity by *in silico* using homology modeled target structure of eEF-2K. In coumarin-chalcone scaffolds, compound 2C was identified as the lead small molecule inhibitor. Initially, we used a rigid protein docking approach^35^ with AutoDock and docking results showed that compound 2C binds to the ATP-binding site of eEF-2K^37-39^ (Supplementary Materials, Fig. S1). In order to take into account, the flexibility of binding pocket residues of model protein^36^ based on MHCK-A when the ligand approaches to the binding site, induced fit docking (IFD) method was considered. IFD results showed that 2C binds similar region identified by rigid docking, however, its binding pose was slightly different, as expected. We then used top-docking IFD pose and performed the long (400-ns) all-atom molecular dynamics (MD) simulations. When we checked the collected trajectories of 2C at the binding site of the target protein, it is found that compound diffuses from the binding pocket (Supplementary Fig. S2). Next, we used top-5 docking poses which all have very similar docking scores changing between (−9.95 to - 8.92 kcal/mol) in MD simulations. For each docking pose, 50-ns all-atom MD simulations are performed, and MD simulations results were shown that poses-2 and 4 did not diffuse from initial docking poses and they were stable (Supplementary Fig. S3). Fig. 1 shows 2D and 3D ligand interactions diagrams of stable docking poses of compound 2C. Throughout the MD simulations, 1000 trajectory frames were collected, and last 500 trajectories were used in free energy calculations of the ligand. The average molecular mechanics generalized Born surface area (MM/GBSA) predicted binding energy scores of poses 2 and 4 of compound 2C were found similar (−57.88 ± 4.20 and -61.86 ± 4.30 kcal/mol, respectively). While Leu137, Met143, Tyr236, Lys238, Asp274 and Gln276 were found in crucial interactions with comound 2C starting the MD simulations with pose-2, corresponding residues were Val168, Lys170, Ile232, Tyr236 and Val278 when we start the MD simulations from pose-4. The main difference between pose-2 and pose-4 throughout the simulations were observed in solvent accessible surface area (SASA) values. Throughout the simulations, initiated with pose-2, we observed water molecules that were bridging the hydrogen bonds between residues and compound 2C. When the induced charge polarization by the active site of the protein environment is considered, quantum mechanics (QM) modeling may give the highest level of docking accuracy. For these reasons, quantum polarized ligand docking (QPLD) is also considered which uses *ab initio* charge calculations for the docking score comparison with the previously studied eEF-2K inhibitors (Table S1). Results showed that compound-2C (−7.691 kcal/mol) has better docking scores than some of the published eEF-2K inhibitors such as non-potent inhibitor A484954 (−5.621 kcal/mol), NH125 (−4.503 kcal/mol), TX-1918 (−5.533 kcal/mol) except non-specific eEF-2K inhibitor Rottlerin (−8.263, kcal/mol).

**Figure 1.**
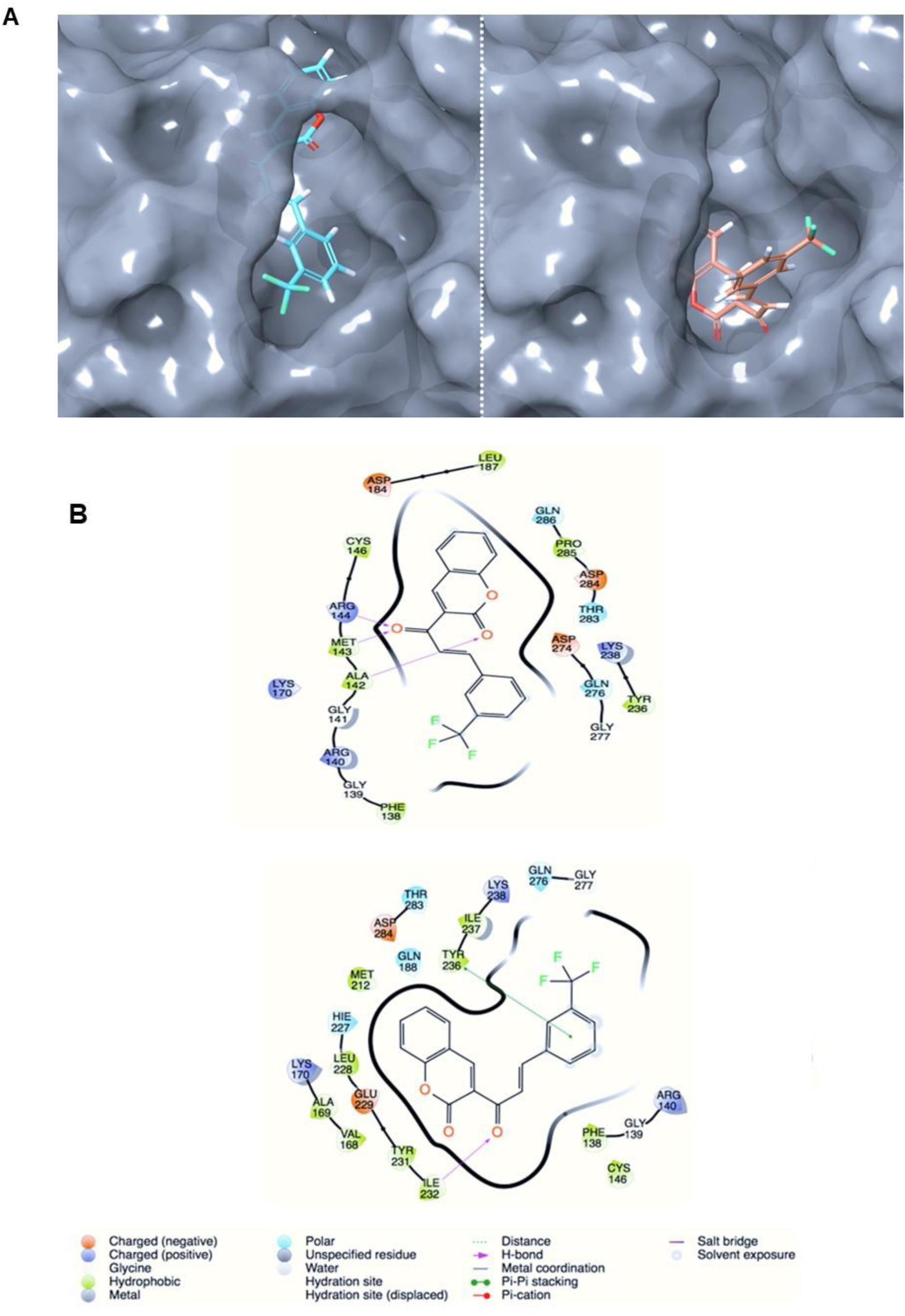
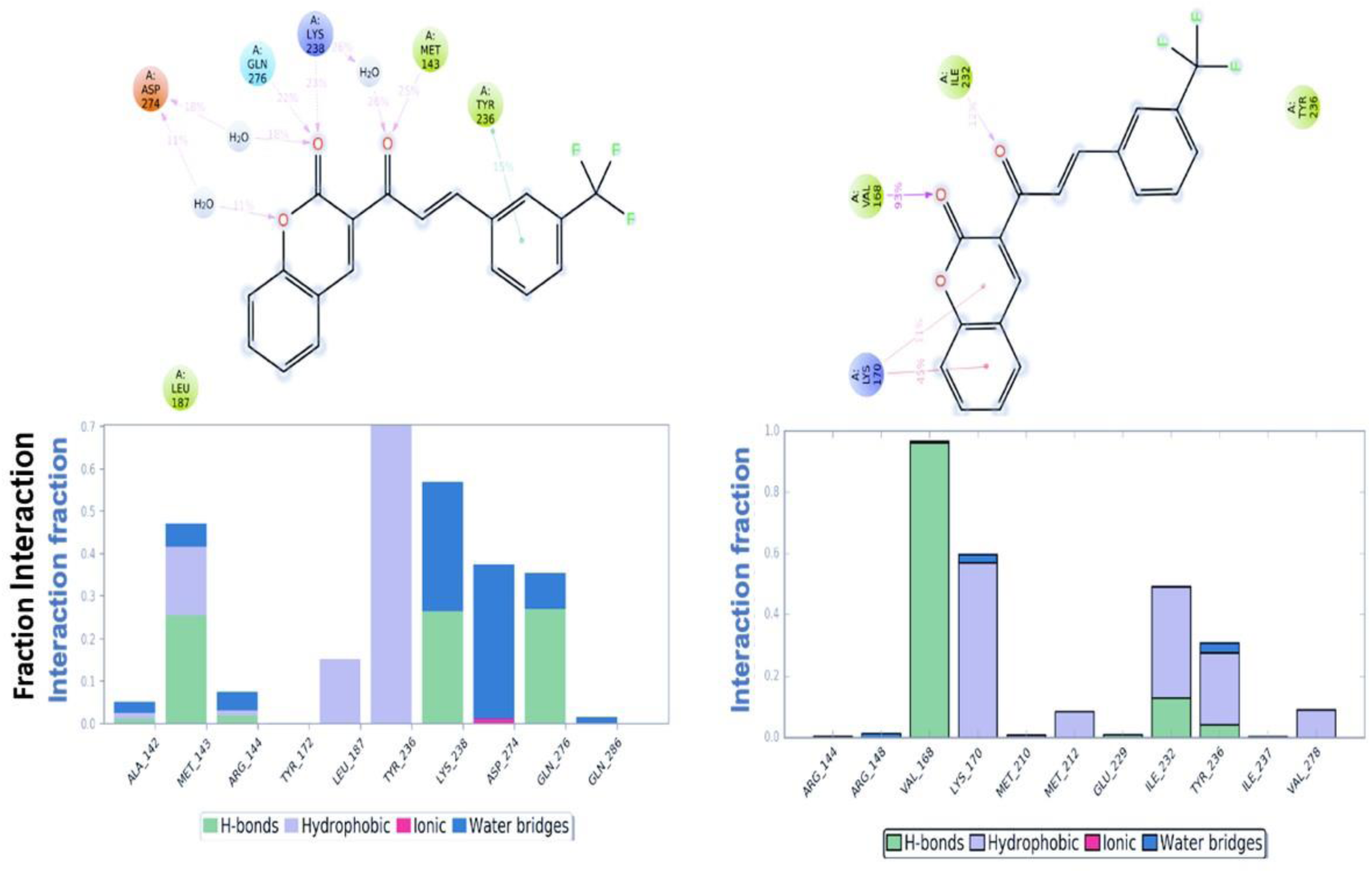
Docking and molecular dynamics interaction sof Compund 2C and EF2K. (a, b) 3D and 2D ligand interactions diagram of **2C** (docking poses); (left) pose-2, right (pose-4). (c) 2D ligand interactions diagram of **2C** (representative MD poses); (left) pose-2, right (pose-4). Figure also shows interaction fractions of crucial residues at the binding.

### Pharmakological properties and predicted ADME profiles of the lead compound 2C

To determine the druglike properties of compound **2C**, we investigated pharmacokinetic properties such as blood brain barrier (BBB, log ratio), lipophilicity, log of compound octanol-water distribution (G-LogP), human serum protein binding (Prot-bind, %), water solubility (WSol, log mg/L), human hERG channel inhibition (hERG-inh, pKi), human serotonin transporter inhibition (SERT-inh, pKi) of compound **2C**^40^ (Supplemnentary data, Table S2). The data for the BBB penetration model is expressed as log values of the ratio of the metabolite concentrations in brain and plasma. According to these calculations, the cutoff value of this parameter should be between -0.3 and 1.5 in MetaCore/MetaDrug tool. Since the calculated BBB value of **2C** is very close to the cutoff value, **2C** is predicted topenetrate through BBB. The lipophilicity value of compound **2C** also shows the permeability capacity. As seen inthe Table S2, human hERG channel inhibition (pKi) value of compound **2C** was determined as - 0.27, suggesting that compound **2C** may not play a role as hERG channel blocker, thus its cardiotoxicity risks (i.e., long QT syndrome) will be limited. However, predicted human serum protein binding percentage basaed on QSAR is found be high, suggesting that compound 2C can achieveeffective drug concentrations and is very informative in establishing safety margins.

### Compound 2C inhibits cell proliferation and colony formation of breast cancer cells

The effect of compound **2C** on cell proliferation and colony formation was evaluated in various breast cancer cell lines, including MCF-7 (Estrogen receptor positive, ER+), and TNBC cells such as MDA-MB-231 (p53 mutated), BT-20 (PI3K and p53mutated) and MDA-MB-436 (BRCA1 mutated)^14^. As shown in Fig. 2a, treatment with the compound **2C** dramatically reduced cell proliferation and colony formation in breast cancer cell lines (Fig. 2a). Compound **2C** inhibited cell proliferation and the number of colonies in at 1 µM or lower concentrations in most of the cell lines compared to DMSO treated control cells while previously published A484954 did not inhibit cell proliferation up to 50 µM drug concentrations in MDA-MB-231, MDA-MB-436 and BT-20 cells (Fig. 2 c-e). We also investigated the effect of **2C** on normal breast epithelial cells. The results showed that compound **2C** did not have any effect on normal immortalized human mammary epithelial cell line (MCF-10A) (Fig. 2e,f).

**Figure 2.**
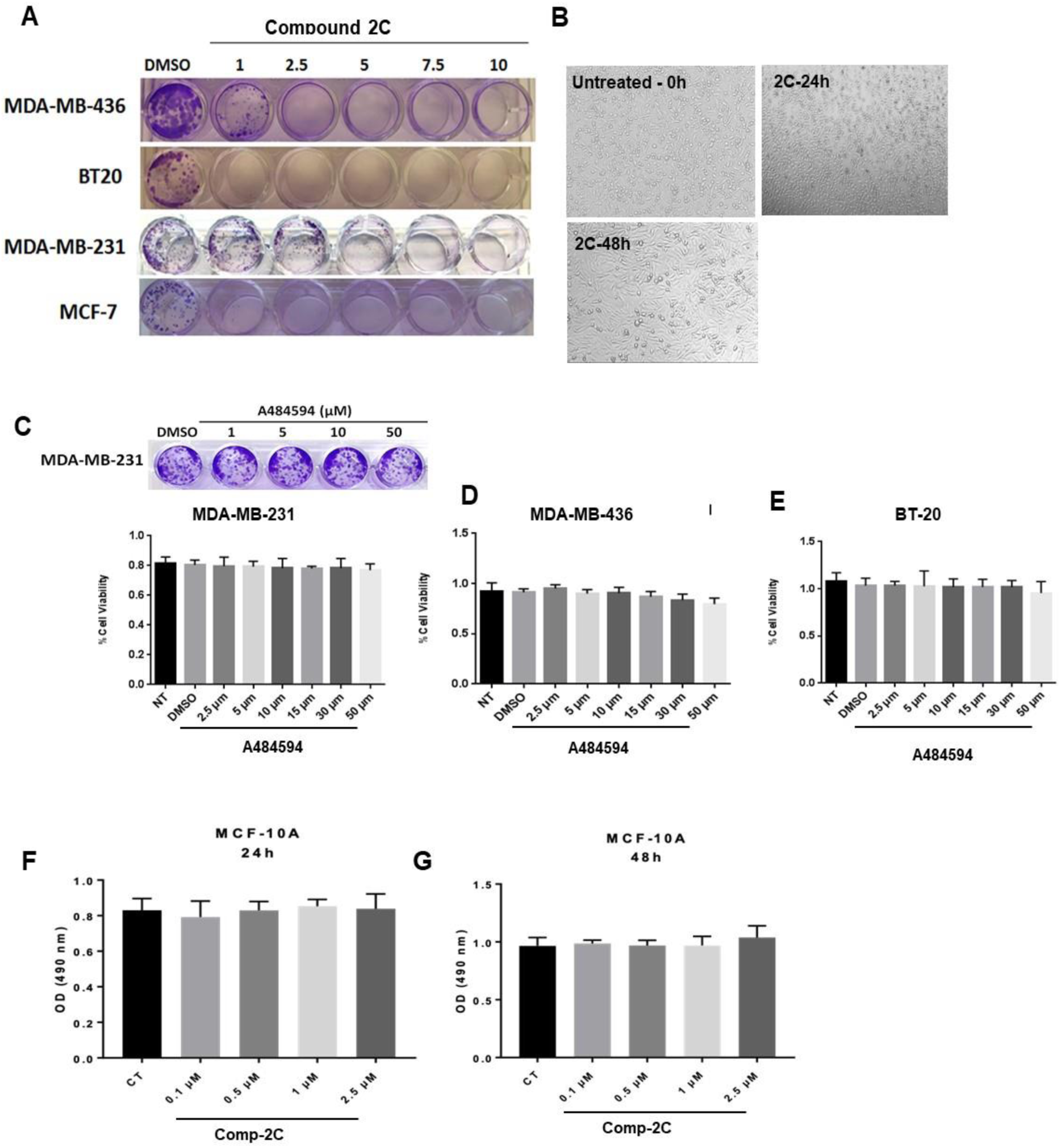
Compound 2C inhibits proliferation and colony formation in breast cancer cells. **a**. Effects of compound **2C** on the colony formation of MCF-7, MDA-MB-436, BT-20 and MDA-MB-231 cells. Compound **2C** inhibits colony formation in the cells and significantly decreased the number of colonies in a dose-dependent manner. MDA-MB-231 cells treated with **2C** cells were imaged by light microscopy at 24h and 48h. **c-e**. Treatment of breast cancer cells with A484954, a previoulsy reported EF2K inhibitor, did not inhibit cell proliferation upto 50uM drug concentrations in MDA-MB-231, MDA-MB-436 and BT-20 cells. **f, g**. Compound **2C** did not show any toxicity on normal immortalized human mammary epithelial cell line for 24h and 48h (MCF-10A). Cell colonies were stained with crystal violet and the number of colonies was quantified after 8–14 days.

### Compound 2C inhibits eEF-2K activity in breast cancer cells

To determine the effect of **2C** on eEF-2K activity in breast cancer cells, we treated cells and evaluated inhibition of eEF-2K by examiming p-EF2 (Thr56) levels by Western blot^13,14^. Treatment with compound **2C** led to a marked inhibition of eEF-2K as indicated by reduced levels of p-EF2, a direct downstream target of eEF-2K, at 1 µM in breast cancer cells up to 48 h. GAPDH and *β*-actin were used as loading controls. Compound **2C** showed strong inhibition against eEF-2K at 0.1 µM (Fig. 3a). Compound **2C** treatment inhibited eEF-2K in in a dose dependent manner at increasing doses (1, 2.5 and 5 µM) at 6h (Fig. 3b). The time-dependent inhibition of eEF-2K by compound **2C** (1 µM) was evaluated in TNBC MDA-MB-231 and ER+MCF-7 cells (Fig. 3c,d). Our results indicated that inhibition of eEF-2K as indicated by reduced p-EF2 (Thr56) levels lasted upto 48 h (Fig. 3c,d).

**Figure 3.**
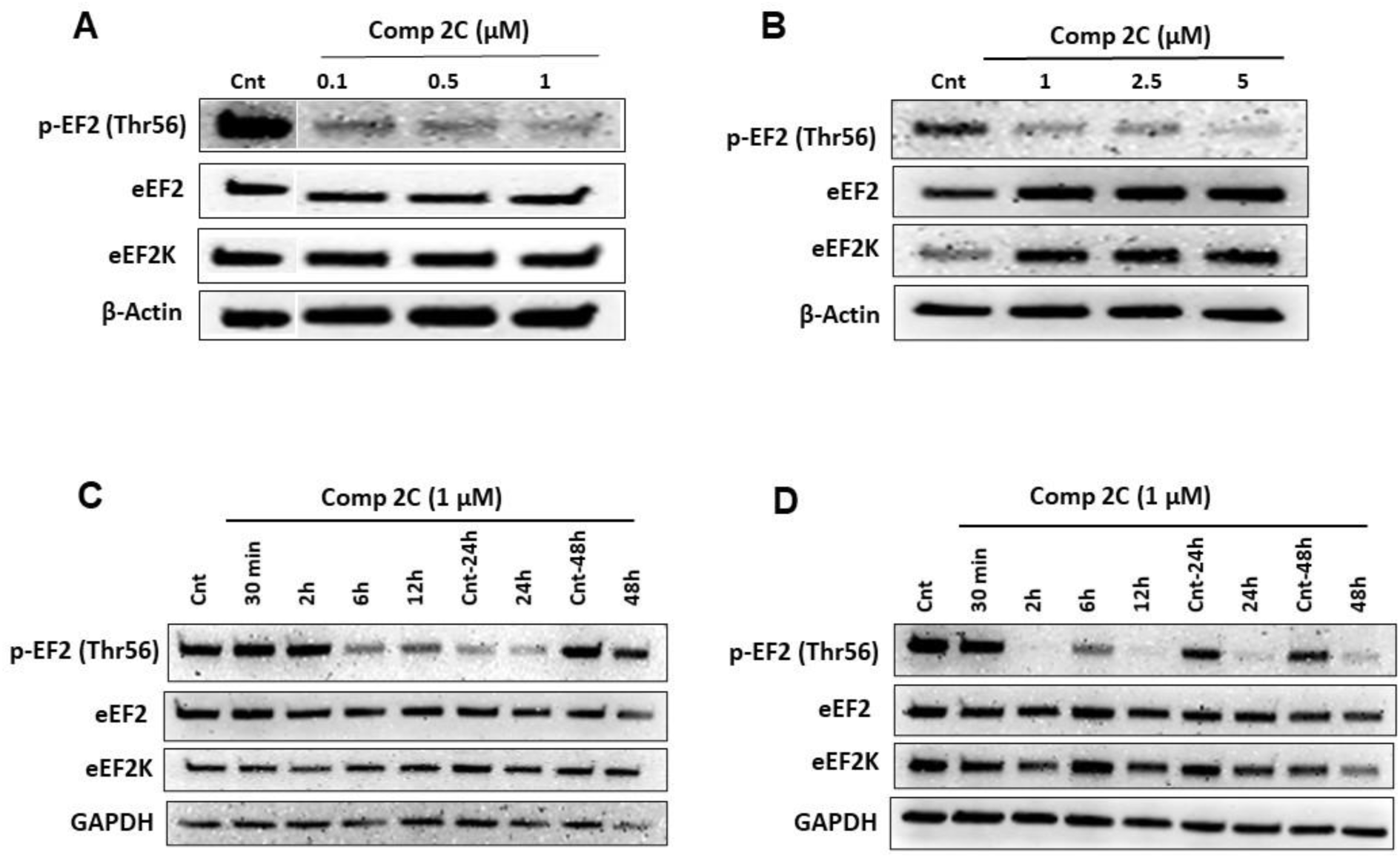
Compound 2C inhibits activity of eEF-2K as indicated by reduced phosphorylation of its downstream target EF2. **a**. MDA-MB-231 cells were treated with compound **2C** in the range of 0.1-1 μM and pEF2 levels analyzed by Western blot. **b**. BT-20 cells were treated with compound **2C** increasing doses of 1 and 5 μM for 6h. **c**. MDA-MB-231 cells were treated with compound **2C** between 0.1 and 1 μM for 2h and p-EF2 levels were evaluated. **d**. MCF-7 cells were treated with compound **2C** at 1 μM upto 48h pEF2 levels analyzed by Western blot.

### Compound 2C induces apoptosis and alters cell cycle distrubition in human breast cancer cells

To evaluate of programmed cell death following the treatment of compound **2C** (2.5, 5 and 10 µM, for 72 h) in breast cancer cells, we first investigated the induction of apoptosis by Annexin V-FITC and PI staining assay staining. These results revealed that compound **2C** induces apoptosis in all breast cancer cell lines. Compound **2C** induced dramatic increase in apoptotic cells in MCF-7 cells, leading to total number of late and early apoptotic cells percentage as high as 91.38 % at the higest dose (Fig. 4a,b). Compound **2C** significantly induced apoptosis in 70.8 % of MDA-MB-231 cells compared to control treated cells (Fig. 5a,b). MDA-MB-436 and BT20 cells underwent apoptotosis by **2C** treatment, however number of apopototic cells much lower compared to MCF7 cells (Fig. 6a,b and 7a,b). To evaluate the mechanism by which **2C** inhibits cell proliferation, breast cancer cells were treated with **2C** and analyzed by cell cycle distribution and flow cytometry. Results showed that significant accumulation in G1 and G2 and reduced SM phases following the treatment of compound **2C** (Fig. 4-7c,d).

**Figure 4.**
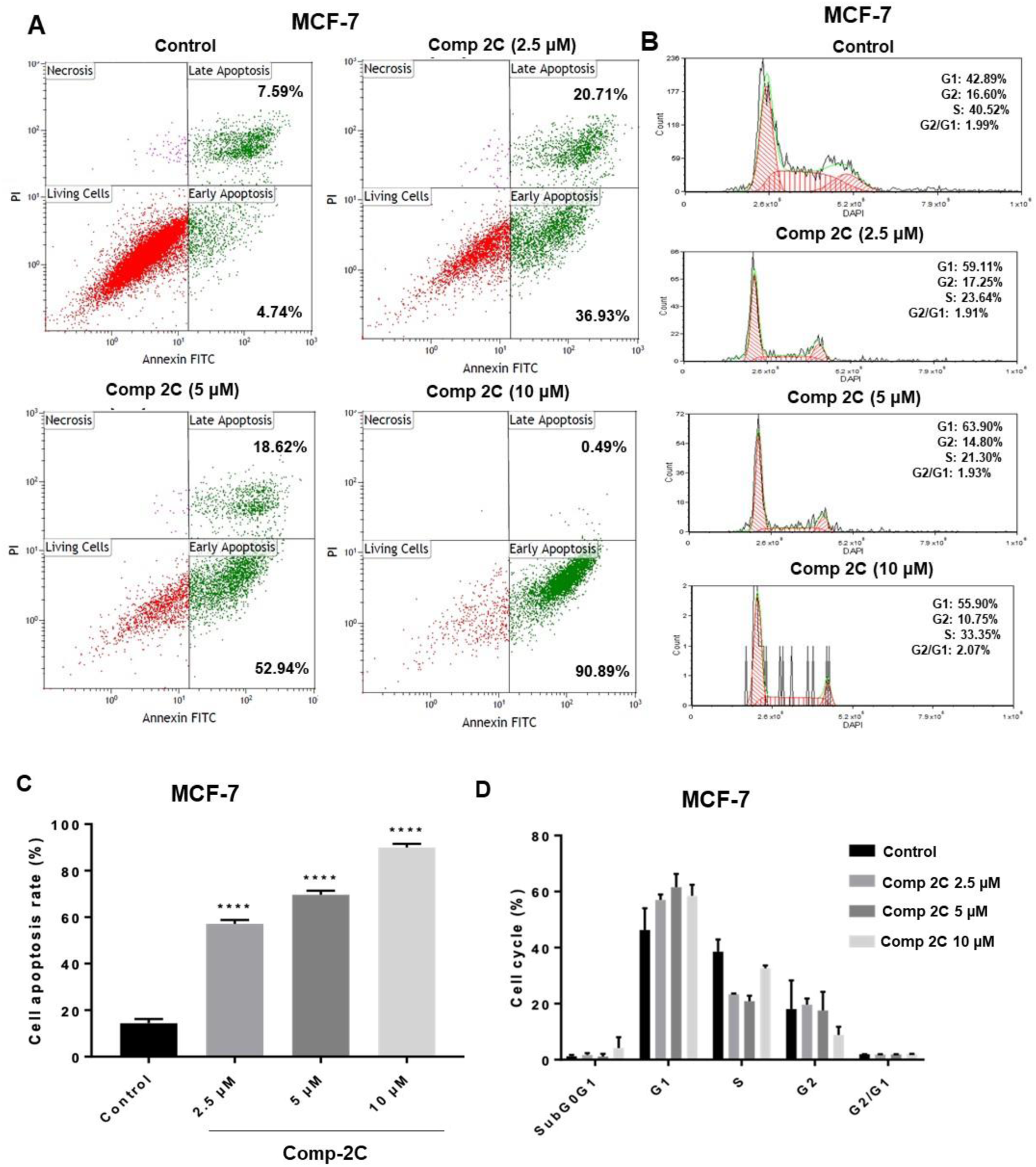
EF2K inhibitor Compound 2C induces apoptosis in ER+ breast cancer cells. Compound **2C** induces apoptosis in MCF-7 cells. **a**. MCF-7 cells were treated with compound **2C** (2.5-10 µM) for 72h or left untreated and analyzed by PI staining and FACS. **b**. Effect of compound **2C** on cell cycle distribution in breast cancer cells. MCF-7 cells were treated with compound **2C** and cell cycle was evaluated by FACS. **c**. and **d**. Bar graphs show the percentages of the cell apoptosis and cell cycle for MCF-7 cells.

**Figure 5.**
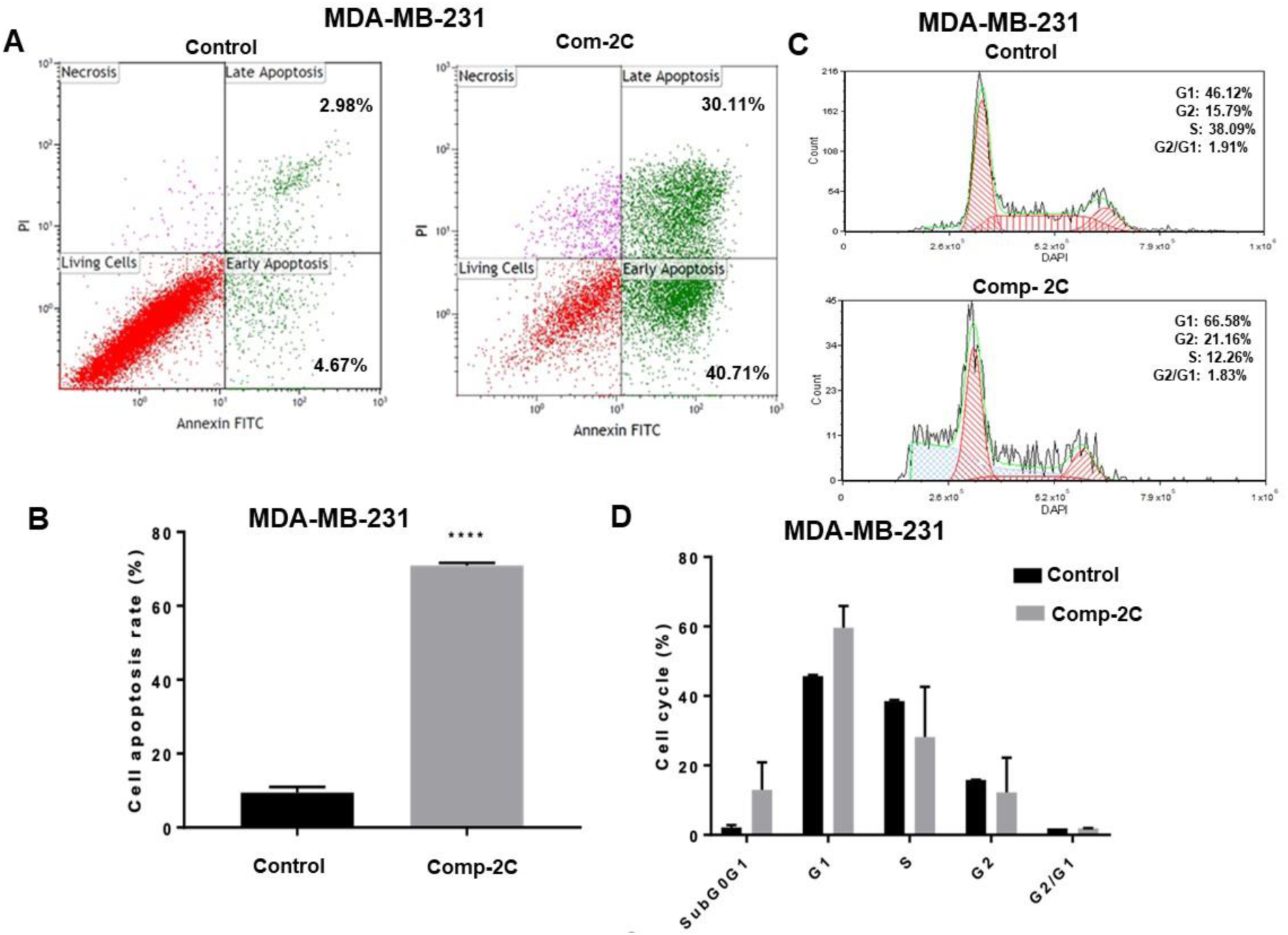
EF2K inhibitor Compound 2C induces apoptosis in TNBC cells. **a**. MDA-MB-231 cells were treated with compound **2C** at 10 µM for 72h. Annexin V-FITC and PI staining assay was used to determine the apoptotic cell death. **b**. The levels of cell apoptosis showed a significant difference in the cells compared to DMSO. **c**. Cell cycle was evaluated by FACS. **d**. Bar graphs show the percentages of cell cycle for MDA-MB-231 cells.

**Figure 6.**
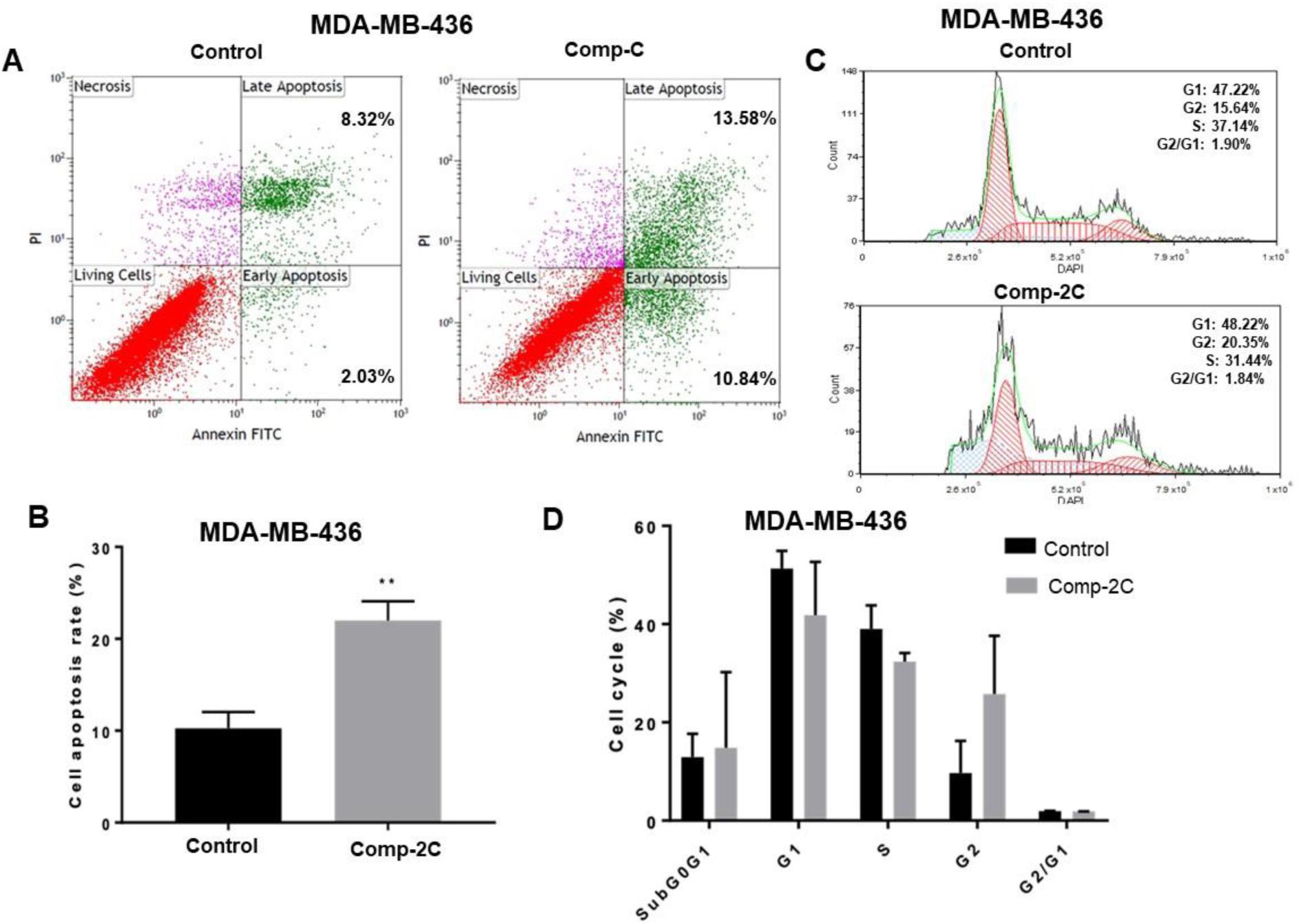
EF2K inhibitor Compound 2C induces apoptosis in TNBC cells. **a**. MDA-MB-436 cells were treated with compound **2C** for 72h. Annexin V-FITC and PI staining assay was used to determine the apoptotic cell death. **b**. The levels of cell apoptosis showed a significant difference in the cells compared to DMSO. **c**. Effect of cell cycle was evaluated by FACS. **d**. Bar graphs show the percentages of cell cycle for MDA-MB-436 cells.

**Figure 7.**
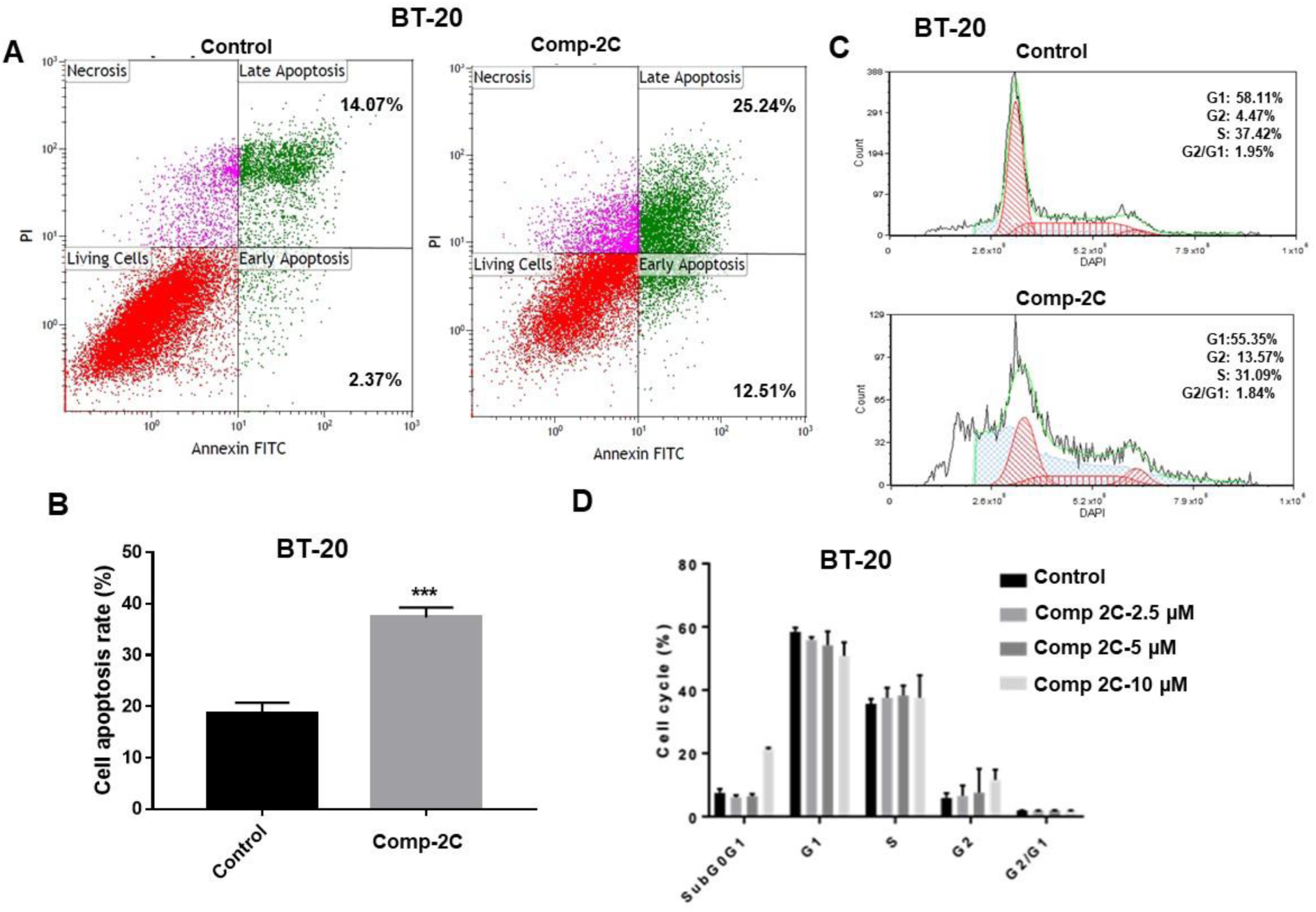
Compound 2C induces apoptosis in BT-20 cells. **a**. BT-20 cells were treated with compound **2C** for 72h. Annexin V-FITC and PI staining assay was used to determine the apoptotic cell death. **b**. The levels of cell apoptosis showed a significant difference in the cells compared to DMSO. **c**. Effect of cell cycle was evaluated by FACS. **d**. Bar graphs show the percentages of cell cycle for BT-20 cells.

### *In vivo* systemic administration of the EF2K inhibitor encapsulated in lipid nanoparticles suppresses growth of orthotopic tumor xenografts in TNBC tumors in mice

To determine the *in vivo* eEF-2K inhibitory effect and therapeutic efficacy of compund **2C** in highly aggressive TNBC tumor models, MDA-MB-231 cells were orthotopically implanted into the mammary fat pad in nude mice. About 2 weeks later, single-lipid nanoparticles incorporating **2C** (5 mice/group) were intraperitoneally (i.p) administered twice a week at 20 mg/kg dose for four weeks. As shown in Fig. 8a, treatment with single lipid nanoparticles incorporating **2C** significantly inhibited tumor growth in mice (p<0.05). Treatment with compound **2C** inhibits for 4 weeks did not lead to any observed toxicity as indicated by behavioral change, appearance and weight loss in mice (Fig 8b). Furthermore, we assessed the role of the inhibitor in inhibiting eEF-2K activity in *in vivo* TNBC tumors. As shown in Fig. 8c eEF-2K inhibitor treatment led to significant reduction in eEF-2K activity as indicated by reduced levels of p-EF2 (Thr56) in MDA-MB-231 tumor xenographs in mice by Western blot analysis compared to control treated tumors.

**Figure 8.**
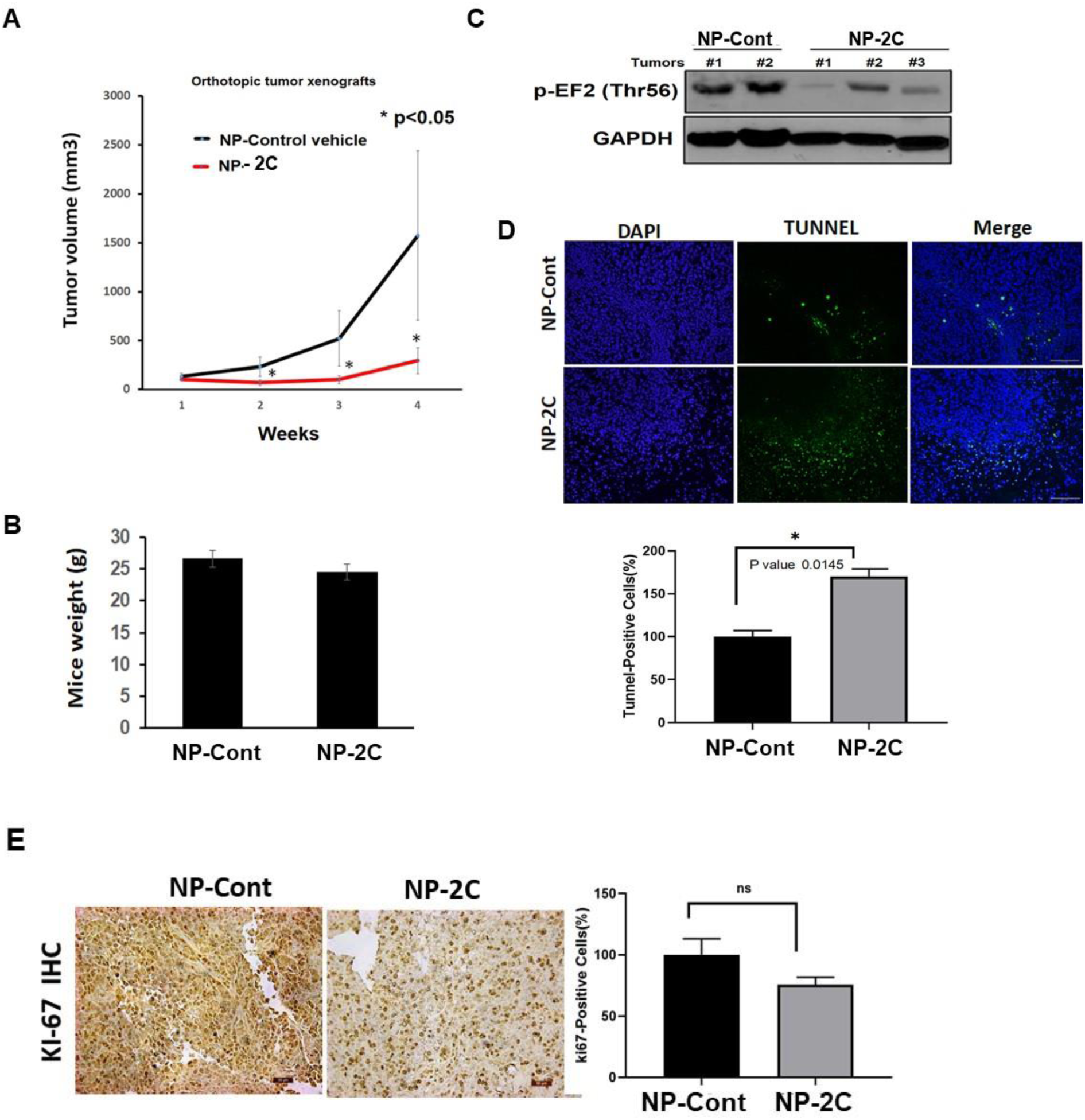
*In vivo* treatment with EF2K inhibitor 2C supressed \growth of highly aggressive TNBC (MDA MB-231) tumors in orthotopic xenograph models in mice. **a**. The mice were sytemicall treated with either single-lipid liposomal nanoparticles (NP) incorporating compound **2C** (20 mg/kg) or empty control nanoparticles. Tumor volumes were measured weekly by electronic calipers and are shown as mean ± SD. b. EF2K inhibitor 2C is safe and did not cause any toxicity in mice. Treatment with EF2K inhibitor **2C** for 4 weeks did not cause any negative impact in eating habits of mice, apprearce of mice, behavioral change, or weight loss and any observed toxicity. **c**. Tumor samples from mice were collected 24 h after the last NP-2C or NP-control (empty vehicle) treatment and analyzed for pEF2 expression by Western blot. GAPDH was used as a loading control. d,e. Immunohistochemical staining was used to evaluate the expression of *in vivo* apoptosis marker TUNEL and proliferation marker Ki-67, microvessel density marker CD31, and in MDA-MB-231 mouse xenografts treated with NP-2C or control NP-mimic. Positively stained cells in both treatment groups were quantified (below panel) (Scale bar = 100 nm).

### EF2K inhibitor 2C is did not cause any toxicity in mice

Treatment with EF2K inhibitor **2C** for 4 weeks did not cause any negative effects in eating habits of mice and behavioral change, apprearce of mice or led to any observed toxicity. As shown in Fig 8b treatment with EF2K inhibitor NP-**2C** for 4 weeks did not cause weight loss in mice, suggesting that EF2K inhibitor 2C is effective and safe for *in vivo* applications.

### NP-2C therapy induces tumor cell apoptosis *in vivo*

Breast tumor cell apoptosis was assessed via immunofluorescent staining for TUNEL in tumors obtained in highly aggressive MDA-MB-231 tumor model. NP-2C-based therapy led to induction of significant apoptostic cell dead in MDA-MB-231 tumors *in vivo* (p=0.014) Fig 8c. Photomicrographs (magnification, 200x) of TUNEL-positive breast tumor cells (green) and tumor-cell nuclei (blue) are shown (Fig. 8d). The expression of Ki-67, an intratumoral proliferation marker, was visualized using IHC in breast tumor xenografts in mice after 4 weeks of treatment with **2C** (Fig. 8e) and quantified using densitometry with mean (± SD) values. Although there was a trend for inhibition of Ki-67 it was not statistically significant.

## DISCUSSION

The findings presented here suggest that compound **2C** is a novel and highly potent inhibitor of eEF-2K with significant in vivo activity. We demonstrated that compound **2C** binds to eEF-2K with high affinity, inhibits its activity at submicromolar concentrations and suppresses breast cancer proliferation. More importantly, *in vivo* treatment with compound **2C** encapsulated in single-lipid nanoparticles significantly supressed growth of highly aggressive triple breast cancer tumor models in mice. Our study provides an *in vivo* effective small molecule eEF-2K inhibitor that may be considered for further development of molecularly targeted threapies in TNBC and other cancers that overexpress eEF-2K.

TNBC is characterized by lack of molecular targets (e.g., ER, PR, and HER2 receptors) and agresive course and realy metastasis. TNBC patients can not benefit from avaiable targeted therapies inculuding anti-estrogen hormone or HER2-targeted antibody threapies. Currently, there is no FDA-approved targeted therapy for precision medicine for TNBC patients and majority of TNBC are treated with the conventional chemotherapies such as anthracyclines (e.g., doxorubicin) and taxane-based therapeutics. Identification of new molecular targets and potent inhibitor of these targets are critical for development of novel targeted treatments in TNBC to improve poor prognosis and dismal patient suvival and reduce high mortality rates. Recently, using genetic methods we have previously validated eEF-2K as a potential molecular target TNBC ^13,14,19^ pancreatic^15,16^ and lung cancer^17^.

Highly potent and effective small molecule inhibitors targeting eEF-2K is greatly needed for clinical translation and targeting cancers that rely of eEF-2K for tumor growth and progression. In the current study, we discovered that coumarin-chalcone compounds may be potent and new inhibitors for eEF-2K and can be used for the purpose of developing clinically applicable targeted therapies. One of the major advantages of coumarins is that are associated with low toxicity^41^. Chalcones are linked by a *α,β*-unsaturated carbonyl system^42^. Interestingly, one of the chalcones (i.e, rottlerin, a natural compound) has been reported act as a non-secific eEF-2K inhibitor and inhibits various other protein kinases at concentrations lower than those needed to inhibit eEF-2K^16,28^. In the present study, we found that the presence of CF_3_ group at 3-position on the phenyl ring of coumarinyl chalcone structure enhanced its eEF-2K inhibitor activity due to its electron withdrawing group properties and inductive effects of trifluoromethyl group.

NH125 has been reported to be the first eEF-2K inhibitor in the literature. However, we and others demonstrated that it acts as a non-specific protein-aggregating agent^29,30^. A-484954 is a weak inhibitor of eEF-2K and effective only concentrations higher than 50-75 µM in cells. Another compound, a pyrido [2,3-*b*]pyrimidine-2,4-dione derivative, has been reported but its not more potent than A-484954, and each only caused partial inhibition of eEF2 phosphorylation at 75 μM in MDA-MB-231 breast cancer cells^43^. Additionally, we have shown that a natural compound thymoquinone (TQ) inhibits eEF-2K and decrease the phosphorylation of EF2 and the expression of eEF-2K in a dose-dependent manner at 5 µM in breast cancer cells^44^. Although, TQ was highly effective in inhibiting eEF2K in *in vivo* tumor models at 20mg/kg doses, it is not a specific inhibitor of eEF-2K and inhibits other targets as well, making it difficult use in the clinical settings.

The efforts for developing potent eEF-2K inhibitors have been hindered by the lack of crystal and a 3-dimensional structure for eEF-2K. However, the development of a homology model of eEF-2K based on the known structures of other members of the α-kinase family such as CMAK has marketly increased rationale drug design and of discovering potent and selective eEF-2K inhibitors^36^. In fact, using the homology modeling of eEF-2K using the homology modeling, recently, we identified coumarin-3-carboxamides (A1 and A2) with morpholine and tert-butyl piperazine-1-carboxylate groups attached to the phenyl ring via the methylene bridge as specific eEF-2K inhibitors at 1 and 2.5 µM doses, respectively^36^. Here, we identified as one of the most effective eEF-2K inhibitors (compound **2C**) that about 100-fold more potent and exerts a higher eEF-2K inhibitors activity compared with the known eEF-2K inhibitors (i.e., A-484954, IC_50_ for cell proliferation = 75 µM)^31,32^. More importantly compound **2C** was very effective in inhibiting tumor growth of highly aggressive MDA-MB-231 TNBC model in mice at 20 mg/kg twice a week injections with no observed toxicity, suggesting that **2C** may have a potential for *in vivo* applications.

The topology of binding pocket region of the protein-ligand complex is largely dependent on conformational changes caused by the bound ligand. Unfortunately, unraveling the X-Ray structure of a protein-ligand complex requires considerable time and investment and mostly it is difficult. Since induced-fit docking approach takes into account the flexibility of both ligand and active site residues, with this method possible binding modes and associated conformational changes of both docked ligand and crucial residues at the binding can be explored. Since the correct partial charges of ligand is crucial in docking, we performed quantum mechanics calculations and used these charges at the docking. The eEF-2K does not have a depth binding pocket, so we used top-5 docking poses of compound **2C** and, we performed all-atom MD simulations for these poses. Results showed that two of the top-five poses (poses 2 and 4) of **2C** have stable conformations and they do not diffuse from the binding pocket and constructs interactions with crucial residues. These two poses have similar binding mode with slight difference in conformation, suggesting that data regarding interaction of **2C** with eEF-2K is highly reliable.

Overall, development of *in vivo* active and potent inhibitors targeting eEF-2K is greatly needed for clinical translation for cancer patients. Our findings indicate that compound **2C** is a potent and *in vivo* active and safe inhibitor of eEF-2K with no observed toxicity and induces significant apoptotic response in breast cancer tumors, suggesting compound **2C** may be a potential therapeutic strategy for molecularly targeted strategies in breast cancer. Compound **2C** may be used as a therapeutic agent not only againt breast also for other cancers such as lung and pancreatic cancers that are sensitive to eEF-2K inhibition. Further studies inclusing extensive toxicology, and PK/PD studies are needed to be performed for further development of this compound and completion of preclinical development.

## MATERIALS AND METHODS

### *In silico* Pharmacokinetic profiles of Compund 2C

To investigate of *in silico* pharmacokinetic profiles of compound **2C**, the MetaCore/MetaDrug platform from Clarivate Analytics was used^40^. MetaCore/MetaDrug uses binary QSAR models for the prediction of the pharmacokinetic properties. The prediction of a therapeutic activity or toxic effect using recursive partitioning algorithm is calculated based on the ChemTree ability to correlate structural descriptors to that property. These pharmacokinetic properties successfully investigated the predicted toxicityand adsorption, distribution, metabolism, excretion (ADME) of compound **2C**.

### Molecular Modeling Studies-Ligand Preparation

Compound **2C** was prepared with the OPLS3 forcefield using LigPrep module (Schrodinger Release 2015-2, LigPrep) of Maestro molecular modeling program. The protonation states at neutral pH 7 was determined by Epik module^45^. Prepared ligands were further optimized by quantum mechanics (QM) method using Jaguar module of Maestro. For this aim, 6-31G* basis set, B3LYP density functional and “ultrafine” SCF accuracy level were used. Mulliken partial charges were recorded and these partial charges were used in further docking studies.

### Protein Preparation

Model 3D structure of protein was used as target. 5DYJ PDB coded protein was used as a template to produce the 3D structure of the target protein. Homology modeling studies were performed with Swiss Model. The target protein was prepared with Protein Preparation module of Maestro. Protonation states of the residues were estimated in PROPKA at blood pH 7.4. Then, the refinement of protein coordinates with geometry optimization using the OPLS3 force field was performed^46^.

### Docking studies were performed with extended induced fit docking (IFD) simulations

As rigid docking, we have used Autodock 4.2. The docking procedure was used a rigid protein^35^ and a flexible ligand were identified with Lamanckian genetic algorithm. The grid was adjusted to points 126, 126, and 126 in x, y and z directions with a grid spacing of 0.375 Å and a distance-dependent function of the dielectric constant was used to calculate the energetic map. For flexible docking, we performed induced fit docking (IFD); thus, initially, Glide/SP was performed, and in geometry optimization, 5.0 Å around the docking poses was used. The refined binding pocket of the target protein^46^ was then used in the redocking procedure using the Glide/XP protocol^47-49^. In QPLD calculations, initially, Glide/SP docking was carried out to generate 10 poses per docked compounds. These poses were submitted to QM charge calculations which uses the 6–31G*/LACVP* basis set, B3LYP density functional, and ‘Ultrafine’ SCF accuracy level.

### Molecular Dynamics (MD) Simulations

The docking poses were used in MD simulations as input coordinates. TIP3P water models used in solvation with 10.0 Å from the edges of protein were used for the determination of the solvation box. For MD simulations, the Desmond program was used. Similar MD protocol to our previously reported studies was used^36^. Last 50% of the trajectories from the MD simulations were used in free energy calculations. For this aim, MM/GBSA calculations from Prime were performed^50,51^.

### Sythesis method

Conventional synthesis method was applied to obtain 3-((E)-3-(3-(trifluoromethyl)phenyl)acryloyl)-2H-chromen-2-one) (compound **2C**) according to the literature as described previously^52,53^. We used 3-acetylcoumarin as a starting material and it was synthesized with a high yield by Knoevenagel condensation reaction of salicylaldehyde and ethylacetoacetate using piperidine in ethanol under reflux^54^. This condensation was used in the second step in order to obtain substituted coumarinyl chalcone in *n*-buthanol as a solvent^52,53^.

### Cell lines and reagents

American Type Culture Collection (ATCC) (Manassas, VA, USA) provided all cell lines including human triple-negative breast cancer (TNBC) cell lines (ER-, PR-, and HER2-) such as MDA-MB-231, MDA-MB-436, and BT-20, ER+ MCF-7. Dulbecco’s Modified Eagle’s Medium (DMEM)/F12 was used for cultuvation of TNBC and MCF-7 cells were with their supplements such as 10% FBS and 1% penicillin streptomycin^45^.

### Cell proliferation and colony formation assay

Colony formation in the cells was detected with compound **2C** and clonogenic assay was performed according to our previous study^36,44^. For this purpose, breast cancer cells (300 cells/well) were seeded in 24-well plates and treated with changing doses of compound **2C** (1, 2.5, 5, 10 μM and at decreased doses (0.1-1 μM)) and cultured for 8–12 days. Crystal violet was used to stain the colonies and quantified using ImageJ software (National Institutes of Health, Bethesda, MD).

### Protein extraction and Western blotting

Western blot analysis was performed according to our previously reported studies^16,44^. Following the treatment of MDA-MB-231 cells with compound **2C**, primary antibodies including p-EF2 (Thr56), eEF2, eEF2K and β-actin (Sigma) and their secondary antibodies were used to detect the expression levels. Time-response manner studies were performed for compound **2C** in MDA-MB-231, MCF-7 cells at 1 µM concentration from changing times 30 minutes to 48 hours compared to DMSO. The most effective p-EF2 inhibition dose was determined. Finally, time-response manner study was applied to compound **2C** in MCF-7 cell lines. Cell signaling technology antibodies were used in this study.

### Analysis of cell death and cell cycle

To determine the programmed cell death, Annexin V assay was used by using reported previously^16^. Cells were seeded in 25 cm^2^ culture flasks (2 x 10^5^ cells/flask). The cells were treated with increasing concentrations of potent compound **2C** (2.5, 5 and 10 µM) in TNBC cells for 72 h and treated with DMSO that was used as a control Annexin V/propidium iodide (PI) staining assay was applied to analyse the cells according to the manufacturer’s protocol (BD Pharmingen FITC–Annexin V kit, San Diego, CA). FACS analysis was performed to detect and quantified positive cells.

### Orthotopic xenograft tumor model of TNBC

MDA-MB-231 cells (2×10^6^ in 20% matrigel) were injected into the mammary fat pad of each mouse and about 2 weeks later, single lipid nanoparticles incorporating the eEF-2K inhibitor compound **2C** was administered twice a week, at two different doses (20 mg/kg) by i.v. injection. Nanoparticles incoprtaing eEF-2K inhibitor were prepared based on our previously published method by us^44^. Tumor volumes were evaluated weekly by electronic calipers. After four weeks of traetment and 24 h after the last injection mice were euthazied under constrant CO_2_ and tumors were removed and kept in -80 °C. Later, tumor tissues were lysed and analyzed by Western blot for evaluation of eEF-2K inhibition.

### Evaluation of apoptosis by TUNEL (TdT-mediated dUTP Nick End Labeling) assay

Apoptotic events after treatment with liposomal siRNA and/or chemotherapy regimens were determined by the TdT-mediated dUTP nick end labeling (TUNEL) assay (Promega, Madison, WI) in tumor sections, according to the manufacturer’s protocol as described previously^13^.

### Immunohistochemistry

Tumor samples of MDA-MB-231 tumors were from the mice and formalin-fixed and paraffin-embedded tumors were sectioned (5 µm) and stained with hematoxylin and eosin. Immunostaining for Ki67 was performed to evaluate intratumoral cell proliferation according to the manufacturer’s protocol. Slides after staining were analyzed under an Eclipse TE200-U microscope (Nikon Instruments Inc., Melville, NY) as previously described^14^.

### Statistical analysis

Data of three independent experiments was given as mean ± SD. Student’s *t* test was used for statistical analysis. *P* values less than 0.05 were considered statistically significant and indicated by asterisk.

## Acknowledgements

The authors are grateful to TUBITAK (Project No: 215S008) for financial support. F.C.O. would like to thank TUBITAK for scholarships (Project No: 215S008 and TUBITAK-BIDEB 2214A program). The University of Texas-MD Anderson Cancer Center Bridge and NIH-NCI-R01CA244344 funds supported NK and BO.

## Author contributions

All authors read and approved the final version of the manuscript.

## Compliance with ethical standards

### Funding

This study was funded by The Scientific and Technological Research Council of Turkey (TUBITAK) (grant number 215S008) and NIH-NCI 1R01CA244344 grants (BO).

## Ethics Statement

Nude athymic female mice were obtained from MD Anderson Cancer Center, Department of Experimental Radiation Oncology Department. All *in vivo* studies were conducted according to the experimental protocol approved by the MD Anderson Institutional Animal Care and Use Committee.

