## Supplementary File for "Target-driven design of a coumarinyl chalcone scaffold based novel EF2 Kinase inhibitor suppresses breast cancer growth *in vivo*"

[1]Department of Experimental Therapeutics, The University of Texas, MD Anderson Cancer Center, 1515 Holcombe Boulevard, Unit 422, Houston, TX 77030, USA, [2]Department of Medical Biology, Çanakkale Onsekiz Mart University, Faculty of Medicine, 17020 Canakkale, Turkey, [3]Department of Chemistry, Natural Products and Drug Research Laboratory, Çanakkale Onsekiz Mart University, Faculty of Science and Arts, 17020 Canakkale, Turkey, [4]DEVA Holding A.S. Cerkezkoy, Tekirdag, Turkey, [5]Izmir Institute of Technology, Department of Chemistry, Bioorganic and Medicinal Chemistry Laboratory, Turkey, [6]Tekirdag Namik Kemal University, Department of Chemistry, Turkey, [7]Gaziantep University, Institute of Health Sciences, Department of Bioinformatics and Computational Biology, 27310, Gaziantep, Turkey, [8]Gaziantep University, Faculty of Arts and Sciences, Department of Chemistry, 27310, Gaziantep, Turkey, [9]Department of Biophysics, School of Medicine, Computational Biology and Molecular Simulations Laboratory, Bahcesehir University, 34734 Istanbul, Turkey, [10]Center for RNA Interference and Non-Coding RNAs, The University of Texas, MD Anderson Cancer Center, Houston, TX, USA

**[#]Patent pending (WO2019/240701)**

**Figures**

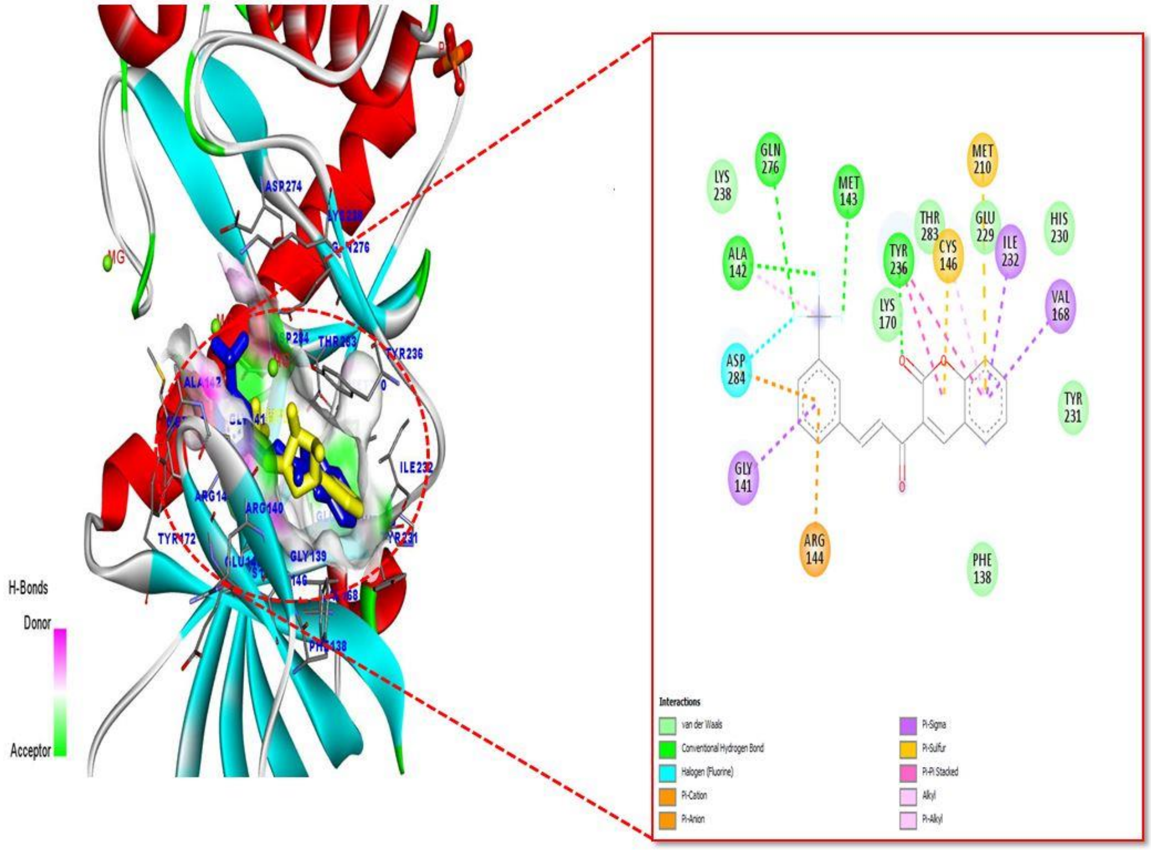

**Figure S1.** The interaction analysis of homology model of eEF-2K with compound **2C**. This compound is bound to ATP binding site of eEF-2K (blue color indicates compound **2C**, yellow color indicates ATP).

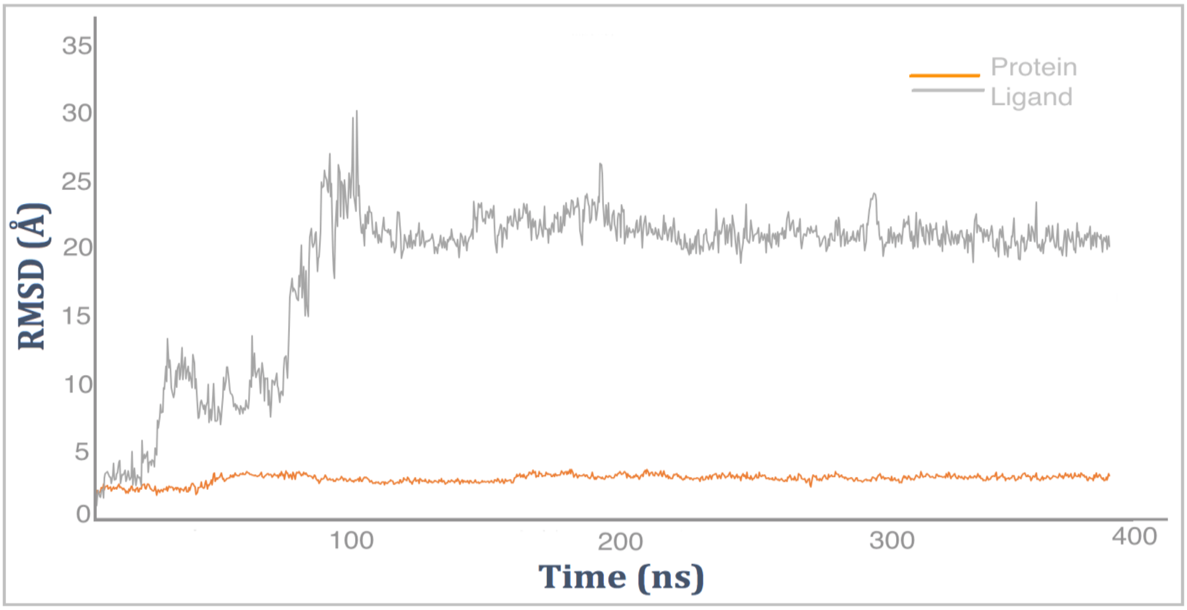


**Figure S2.** RMSD-time plot of protein and top-pose of **2C** RMSDs (based on protein structure) throughout the simulations.

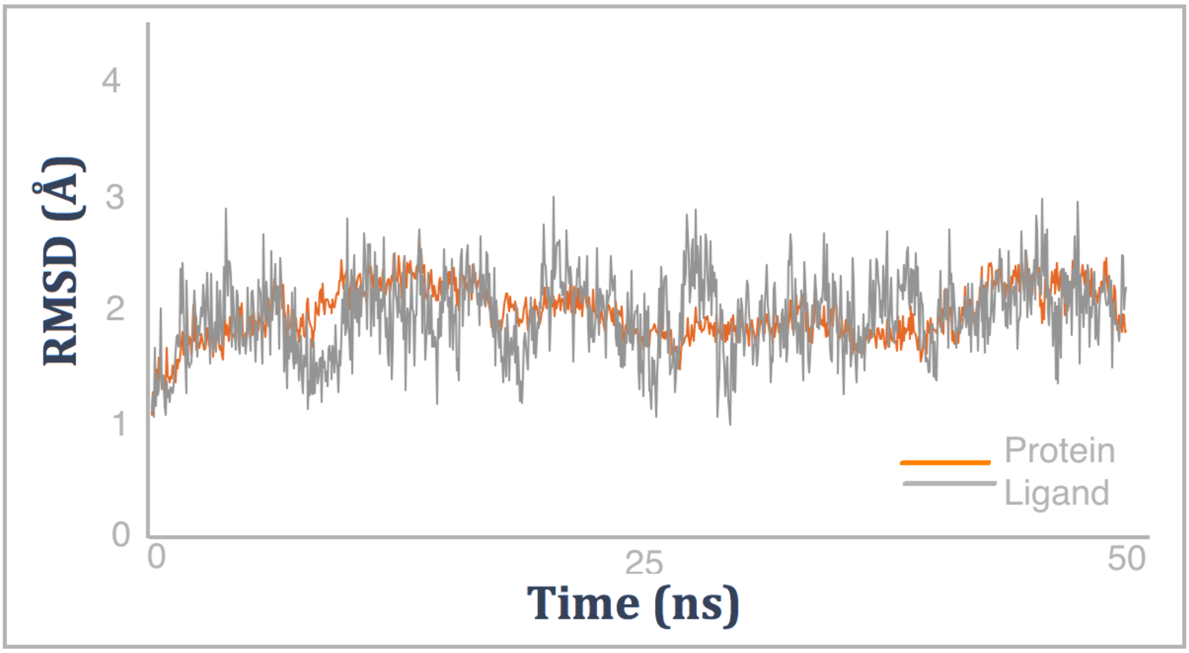


**Figure S3.** RMSD-time plot of protein and second-top-pose of **2C** RMSDs (based on protein structure) throughout the simulations.

| eEF-2K Inhibitors | QPLD scores (kcal/mol) |
|---|---|
| A484954 | -5.621 |
| NH125 | -4.503 |
| TX-1918 | -5.533 |
| ROTTLERIN | -8.263 |
| **Compound 2C** | **-7.691** |

**Table S1.** QPLD docking score comparison of 2C with other eEF-2K inhibitors.

| Compound | BBB, log ratio [1] | G-LogP [2] | Prot-bind, % [3] | WSol, log mg/L [4] | hERG-inh, pKi [5] | SERT-inh, pKi [6] |
|---|---|---|---|---|---|---|
| **2C** | -0.52 | 3.51 | 82.65 | 0.88 | -0.27 | 0.27 |

**Table S2.** Predicted ADME profiles of the compound **2C**

(1) Blood brain barrier penetration model. The data is expressed as log values of the ratio of the metabolite concentrations in brain and plasma. Cutoff is -0.3. Larger values indicate that the metabolite is more likely to enter the brain. Reference: Thomson Reuters. Model description: N=107, R2=0.89, RMSE=0.26.

(2) Lipophilicity, log of compound octanol-water distribution. Cutoffs are -0.4 to 5.6. Values greater than 5.6 correspond to overly hydrophobic compounds. Reference: Syracuse Research, PHYSPROP database. Model description: N=13474, R2=0.95, RMSE=0.21.

(3) Human serum protein binding, %. Cutoff is 50%. A value of more than 95% is highly bound, less than 50% is a low binding metabolite. Reference: Thummel and Shen, 2001 in Goodman & Gilman's The Pharmacological Basis of Therapeutics. Model description: N=265, R2=0.909, RMSE=10.11.

(4) Water solubility at 25 ºC, log mg/L. Cutoffs are from 2 to 4. An acceptable level of solubility is project dependent. Reference: Syracuse Research, PHYSPROP database. Model description: N=2871, 2=0.91, RMSE=0.54.

(5) Human hERG (human ether a-go-go-related gene) channel inhibition , pKi (uM). Cutoff is -1.7. The higher the value, the higher the inhibition activity. Lower values are preferable. Reference: Thomson Reuters. Model description: N=196, R2=0.93, RMSE=0.23.

(6) Human serotonin transporter inhibition, pKi (uM). Cutoff is -1.7. The higher the value, the higher the inhibition activity of the metabolite. Thomson Reuters. Model description: N=256, R2=0.91, RMSE=0.36.
